# Mutation bias shapes the spectrum of adaptive substitutions

**DOI:** 10.1101/2021.04.14.438663

**Authors:** Alejandro V. Cano, Hana Rozhoňová, Arlin Stoltzfus, David M. McCandlish, Joshua L. Payne

**Author notes:** These authors contributed equally (David M. McCandlish, Joshua L. Payne).

## Abstract

Evolutionary adaptation often occurs via the fixation of beneficial point mutations, but different types of mutation may differ in their relative frequencies within the collection of substitutions contributing to adaptation in any given species. Recent studies have established that this spectrum of adaptive substitutions is enriched for classes of mutations that occur at higher rates. Yet, little is known at a quantitative level about the precise extent of this enrichment, or its dependence on other factors such as the beneficial mutation supply or demographic conditions. Here we address the extent to which the mutation spectrum shapes the spectrum of adaptive amino acid substitutions by applying a codon-based negative binomial regression model to three large data sets that include thousands of amino acid changes identified in natural and experimental adaptation in *S. cerevisiae*, *E. coli*, and *M. tuberculosis*. We find that the mutation spectrum has a strong and roughly proportional influence on the spectrum of adaptive substitutions in all three species. In fact, we find that by inferring the mutation rates that best explain the spectrum of adaptive substitutions, we can accurately recover species-specific mutational spectra obtained via mutation accumulation experiments. We complement this empirical analysis with simulations to determine the factors that influence how closely the spectrum of adaptive substitutions mirrors the spectrum of amino acid variants introduced by mutation, and find that the predictive power of mutation depends on multiple factors including population size and the breadth of the mutational target for adaptation.

**SIGNIFICANCE STATEMENT:** How do mutational biases influence the process of adaptation? Classical neo-Darwinian thinking assumes that selection alone determines the course of adaptation from abundant pre-existing variation. Yet, theoretical work shows that under some circumstances the mutation rate to a given variant may have a strong impact on the probability of that variant contributing to adaptation. Here we introduce a statistical approach to analyzing how mutation shapes protein sequence adaptation, and show that the mutation spectrum has a proportional influence on the changes fixed in adaptation observed in three large data sets. We also show via computer simulations that a variety of factors can influence how closely the spectrum of adaptive substitutions mirrors the spectrum of variants introduced by mutation.

## INTRODUCTION

A systematic empirical picture of the spectrum of adaptive substitutions is beginning to emerge from methods of identifying and verifying individual adaptive changes at the molecular level. The most familiar method is the retrospective analysis of adaptive species differences, often in cases where multiple substitutions target the same protein, e.g., changes to photoreceptors involved in spectral tuning [1], changes to ATPase involved in cardiac glycoside resistance [2], or changes to hemoglobin involved in altitude adaptation [3]. Other retrospective analyses focus on cases of recent local adaptation, such as the repeated emergence of antibiotic-resistant bacteria [4, 5] or herbicide-resistant plants [6]. In addition, experimental studies of adaptation in the laboratory provide large and systematic sets of data on the spectrum of adaptive substitutions [7, 8]. While the first two types of studies tend to focus on specific target genes, the third approach, combined with genome sequencing, casts a much broader net, covering the entire genome. Such data were rare just 15 years ago, but they are now sufficiently abundant—cataloging thousands of adaptive events—that accounting for the species-specific spectrum of adaptive substitutions represents an important challenge.

One aspect of this challenge is to understand the role of mutation in shaping the spectrum of adaptive substitutions. Systematic studies of the distribution of mutational types in diverse organisms [9–17] have demonstrated the presence of a variety of biases, including transition bias and GC:AT bias, as well as CpG bias and other context effects (for review, see [18]). At the same time, multiple studies have now shown that adaptive substitutions are enriched for these mutationally likely changes [5, 19–26]. For instance, the influence of a mutational bias favoring transitions is evident in the evolution of antibiotic resistance in *Mycobacterium tuberculosis* [5]. Likewise, the evolution of increased oxygen-affinity in hemoglobins of high-altitude birds shows a tendency to occur at CpG hotspots [24].

Such studies have shown effects of specific types of mutation bias using statistical tests for asymmetry, i.e., tests for a significant excess of a mutationally favored type, relative to a null expectation of parity. A more general question is how strongly the entire mutation spectrum shapes the spectrum of adaptive substitutions. That is, the entire mutation spectrum reflects (simultaneously) all relevant mutation biases, and this spectrum shapes the spectrum of adaptive substitutions to some degree that is, in principle, quantifiable and measurable.

Here, we provide an approach to this more general question, based on modeling the spectrum of missense mutations underlying adaptation as a function of the nucleotide mutation spectrum. More specifically, we use negative binomial regression to model observed numbers of adaptive codon-to-amino acid changes as a function of codon frequencies and per-nucleotide mutation rates, which we derive from experimental measurements of mutation spectra in the absence of selection. This modeling framework allows us to measure the influence of mutation bias on adaptive evolution in terms of the regression coefficient associated with the mutation spectrum.

We separately apply this approach to three data sets of missense changes associated with adaptation in *Saccharomyces cerevisiae*, *Escherichia coli*, and *Mycobacterium tuberculosis*. We find that, in each case, the regression on the mutation spectrum is significant, with a regression coefficient close to 1 (proportional effect) and significantly different from zero (no effect). The ability to predict the spectrum of adaptive substitutions differs substantially amongst the three species, but in each case, we find that experimentally determined mutation spectra provide better model fits than the vast majority of randomized mutation spectra, confirming the relevance of empirical mutation spectra outside of the controlled conditions in which they are typically measured. Moreover, we show that by inferring the optimal mutational spectrum based on the spectrum of adaptive substitutions we can accurately recover species-specific patterns of mutational bias previously documented via mutation accumulation experiments or patterns of neutral diversity. Finally, we use simulations of a population model to explore the possible reasons for differences in predictability of the spectrum of adaptive substitutions. As expected, the impact of the mutation spectrum decreases as the total mutation supply (*Nμ*) increases. However, other factors are important, such as the size and heterogeneity (in adaptive value) of the set of adaptive mutations.

## RESULTS

### Data and model

We curated a list of missense mutations associated with adaptation for each of three species: *S. cerevisiae*, *E. coli*, and *M. tuberculosis* (Fig. 1a,b; Methods). For *S. cerevisiae*, the mutations were associated with adaptation to one of several environments during laboratory evolution, including high salinity [27], low glucose [27], rich media [28], as well as the genetic stress of gene knockout [29]; for *E. coli*, the mutations were associated with adaptation to temperature stress during laboratory evolution [8]; for *M. tuberculosis*, the mutations were associated with natural adaptation to one or more of eleven antibiotics or antibiotic classes, and were derived from clinical isolates [5].

**Fig. 1.**
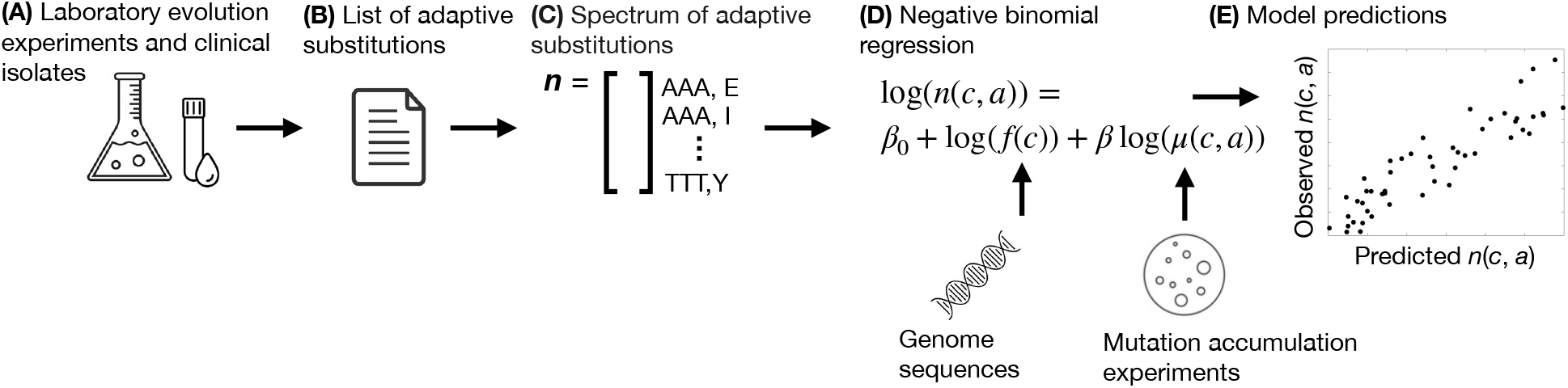
Workflow. (a) We use data from laboratory evolution experiments (*E. coli* and *S. cerevisiae*) and clinical isolates (*M. tuberculosis*) to curate (b) a list of genetic changes associated with adaptation for each species. (c) From each list of adaptive mutations, we construct the spectrum of adaptive substitutions **n**. Each element in this spectrum **n**(*c, a*) corresponds to one of the 354 distinct changes from codon *c* to amino acid *a* that can be produced by a single nucleotide substitution under the standard genetic code and tallies the number of adaptive events per codon-to-amino acid change. (d) We perform negative binomial regression to model the influence of mutation bias on the spectrum of adaptive events, using codon frequencies derived from genome sequences and experimentally characterized mutation spectra. (e) We use the fitted model to predict the spectrum of adaptive events.

Because of the possibility that the same substitutions underlie adaptation in multiple independent populations, we follow [23] in distinguishing between adaptive *paths* defined by a genomic position and a specific mutational change, and the number of substitutional *events* that have occurred along that path in independent populations. For example, the mutational path defined by a G→C transversion in the second position of codon 315 of KatG in *M. tuberculosis*, which changes Ser (AGC) to Thr (ACC), is known to confer resistance to the antibiotic isoniazid [30]. Events along this mutational path are common in adaptation, occurring 766 independent times in our data set. Below, when we construct the spectrum of adaptive substitutions, the data are further aggregated by the *type* of path, out of the 354 possible codon-to-amino-acid paths. For instance, all G→C transversions changing Lys (AAG) to Asn (AAC), at all positions in all genes for a given species, are counted together in the AAG to Asn category for that species, and this same category also includes all G→T transversions that change Lys (AAG) to Asn (AAT). Most codon-to-amino-acid paths, however, include only a single type of nucleotide change, e.g., the Ser (AGC) to Thr path only includes G→C transversions from AGC to ACC as in the KatG example above.

Table 1 reports the number of mutational paths and adaptive events for each of our three species. While the *M. tuberculosis* data set is likely composed solely of adaptive changes (since all mutations included have been experimentally verified to confer antibiotic resistance, [5]), for *S. cerevisiae* and *E. coli*, we expect these data sets to be contaminated with a minority of hitchhikers, i.e., mutations that are not drivers of adaptation but which reached a high frequency due to linkage with a driver. Below, we first present our results under the assumption that the mutations in each data set are exclusively adaptive and then use simulations to assess the robustness of our conclusions to various degrees of contamination.

**TABLE 1.** Data and model fits. Shown are the observed numbers of paths and events for adaptive changes in the three data sets, along with calculated values for the mutation coefficient *β* (with standard error) and its *p*-value, the Pearson’s correlation between observed and predicted spectra of adaptive substitutions and its *p*-value, the number of non-zero elements in the spectrum of adaptive substitutions (out of 354), and the entropy of the spectrum of adaptive substitutions.

|  | Data |  | Neg. binomial regression |  | Prediction model |  | Spectrum elements |  |
| --- | --- | --- | --- | --- | --- | --- | --- | --- |
| Species | Paths | Events | $\beta$ | $p_\beta$ | Correlation | $p_{\text{corr}}$ | Non-zero | Entropy |
| *S. cerevisiae* | 534 | 721 | $1.05 \pm 0.08$ | $< 10^{-16}$ | 0.68 | $< 10^{-16}$ | 265 | 0.91 |
| *E. coli* | 492 | 602 | $0.98 \pm 0.14$ | $< 10^{-11}$ | 0.41 | $< 10^{-14}$ | 176 | 0.80 |
| *M. tuberculosis* | 283 | 4413 | $0.87 \pm 0.23$ | $< 10^{-3}$ | 0.16 | 0.003 | 111 | 0.53 |

For each species, we use the corresponding list of adaptive events to construct the spectrum of adaptive substitutions (Fig. 1c), which we represent as a 354-element vector **n**, where each element **n**(*c, a*) corresponds to a single-nucleotide change from codon *c* to amino acid *a* allowed by the standard genetic code (Methods). For a given species, the value assigned to an element (codon-to-amino acid change) in the spectrum of adaptive substitutions is the observed number of adaptive events associated with that change.

Our goal is to assess the extent to which the spectrum of adaptive substitutions is shaped by the spectrum of genetic changes introduced by mutation (Fig. 1d). To do so, we model the expected number 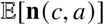 of adaptive mutations from codon *c* to amino acid *a* as being directly proportional to the genomic frequency *f* (*c*) of codon *c*, as well as potentially proportional to the total mutation rate *μ*(*c, a*) of codon *c* to codons for amino acid *a*. We obtained codon frequencies from genomic sequences, and we obtained mutation rates from mutation accumulation experiments and single-nucleotide polymorphism data (Methods). In particular, our model can be expressed as

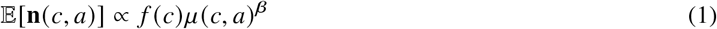

where *β* is an unknown coefficient that describes the dependence of 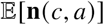 on *μ*(*c, a*). Taking the logarithm of this equation gives

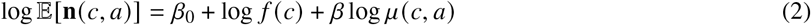

where *β*_0_ determines the constant of proportionality. We use negative binomial regression to estimate *β*_0_ and *β*, which is appropriate for counts data that exhibit over-dispersion [31], such as the data studied here.

Importantly, the regression coefficient *β* in Eqn. 2 measures the influence of mutation bias on adaptation. When *β* = 0, 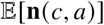 no longer depends on *μ*(*c, a*), implying that mutation bias has no influence on the course of adaptation. When *β* = 1, 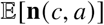 is directly proportional to *μ*(*c, a*), implying a strong influence of mutation bias on adaptation. For instance, *β* = 1 implies that doubling the rate of a particular mutation type doubles the rate of adaptive substitutions of that type. Values of *β* between 0 and 1 indicate an intermediate influence of mutation bias on adaptation. In what follows, we therefore focus on estimating *β* for each of our three species of interest.

### Mutation bias influences adaptation in three distinct species

To what extent does the spectrum of nucleotide changes introduced by mutations influence the genetic basis of adaptive evolution? The three species examined here differ substantially in their mutational spectra (Fig. S1a). *M. tuberculosis* shows the greatest heterogeneity in its mutational spectrum with a 14.5-fold difference between maximum and minimum mutation rates, whereas *S. cerevisiae* and *E. coli* have a somewhat smaller range of rates (5.6-fold and 4.7-fold ranges, respectively). The species also differ substantially in the rates of individual types of nucleotide mutations. For instance the rate of G→C transversion is 2.1-fold higher in *S. cerevisiae* than in *E. coli* (Fig. S1b), whereas the rate of A→T transversions is 2.6-fold higher in *S. cerevisiae* (Fig. S1c) and 3-fold higher in *E. coli* (Fig. S1d) than in *M. tuberculosis*. Simply comparing these mutational spectra to the spectra of adaptive substitutions observed in each species reveals a striking congruence between the rate that different types of nucleotide mutations arise in each species and the frequency that each type of mutation is used in the course of adaptation (Fig. 2a-c).

**Fig. 2.**
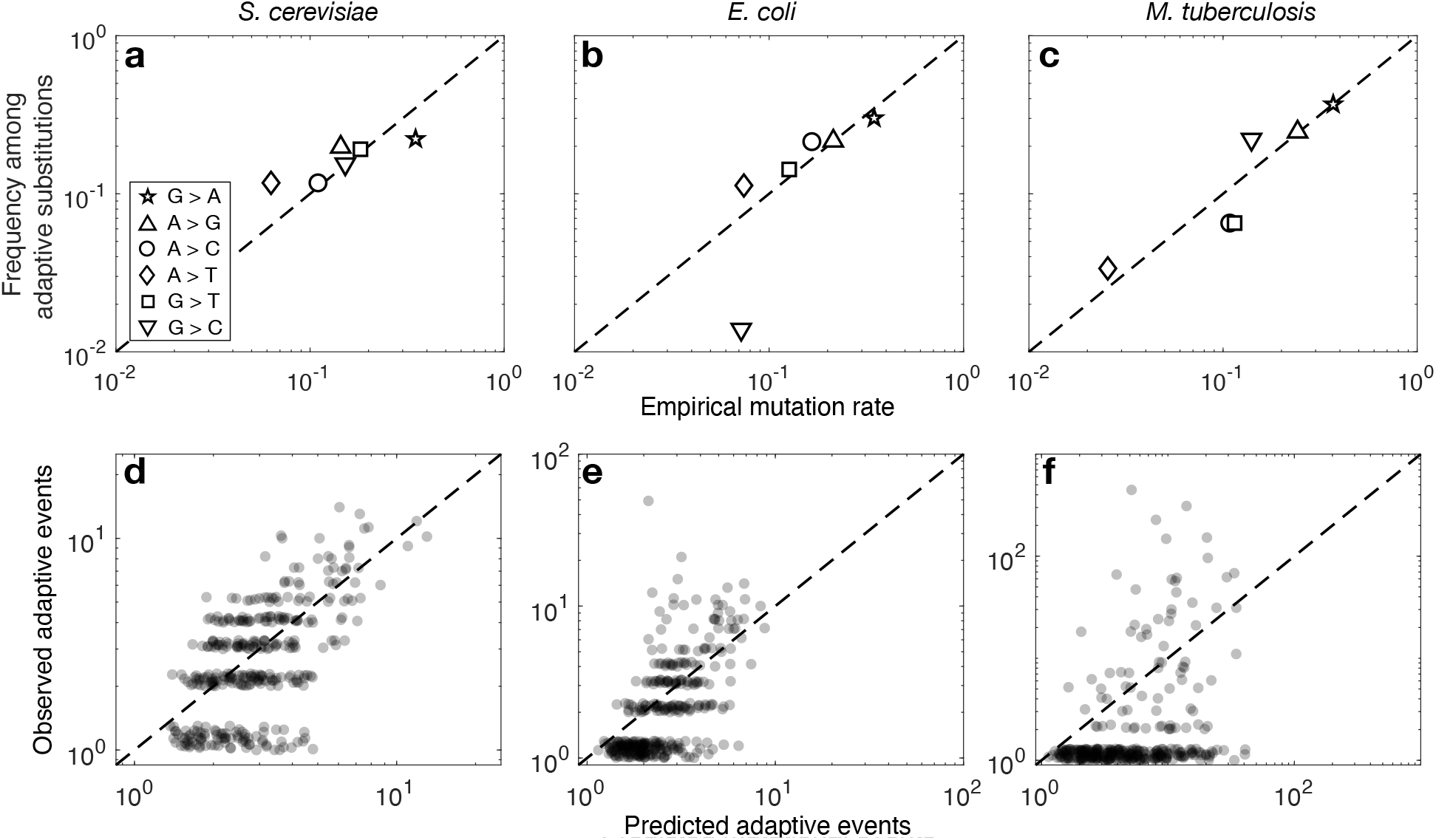
Predicted and observed substitutions at the nucleotide and codon-to-amino acid levels. (a-c) The frequency of nucleotide changes among adaptive substitutions is plotted as a function of the empirical mutation rate for (a) *S. cerevisiae*, (b) *E. coli*, and (c) *M. tuberculosis*. The symbols correspond to the six different types of point mutations (inset in panel a). (d-f) The predicted spectra of adaptive substitutions are shown in relation to the observed spectra of adaptive substitutions for (d) *S. cerevisiae*, (e) *E. coli*, and (f) *M. tuberculosis*. For visualisation purposes, a pseudo count of 1 event and a jitter of range [0,0.3] were added to both the observed and predicted numbers of events in panels (d-f).

While intriguing, the above analysis does not account for the potentially confounding effects of the genetic code and codon usage among the three species, where in particular the three species differ substantially in their patterns of codon usage (Fig. S1e-g). For example GAA (Glu) is the most frequent codon in *S. cerevisiae* (frequency 0.045) and the 2nd most frequent codon in *E. coli* (frequency 0.039), but it appears much less frequently in *M. tuberculosis* (frequency 0.016). Thus, we might expect adaptive GAA→AAA (Glu→Lys) changes to occur more frequently in *S. cerevisiae* and *E. coli* than in *M. tuberculosis*, merely by merit of the greater frequency of GAA in the former two species. To account for this type of influence, as well as for the fact that identical amino acid substitutions can be produced by different nucleotide mutations because of the standard genetic code, we fit a codon-based negative binomial regression model to ask to what extent the mutation spectrum influences the spectrum of adaptive substitutions (Eqn. 2). For each of the three species, this analysis produced an estimate of the regression coefficient *β* that captures the influence of the mutational spectrum on the spectrum of adaptive substitutions, as well as an associated *p*-value under the null hypothesis that mutational biases have no influence on the spectrum of adaptive substitutions (i.e., *β* = 0).

The results, shown in Table 1, reveal a strong and statistically significant influence of mutation bias on adaptation in all three species, with each of the 95 % confidence intervals containing *β* = 1, and excluding *β* = 0. Specifically, for *S. cerevisiae*, *β* = 1.05 (95 % CI, 0.89 to 1.21), for *E. coli β* = 0.98 (95 % CI, 0.71 to 1.25), and for *M. tuberculosis*, *β* = 0.87 (95 %, 0.42 to 1.32), so that in all three species differences in mutation rates produce approximately proportional changes in the spectrum of adaptive substitutions.

Having seen the influence of the mutation spectrum on the spectrum of adaptive substitutions, we can also ask to what extent the mutational spectrum, pattern of codon usage, and the structure of the genetic code are jointly sufficient to explain the spectrum of adaptive codon-to-amino acid changes observed in each species. In particular, Figure 2d-f shows the observed frequency of each type of codon-to-amino acid change in relation to its predicted frequency under our fitted models. We observe from this figure that despite the mutational spectrum having its maximum theoretically predicted influence (*β* ≈ 1), the predictive power of our model nonetheless differs substantially among the three species, with a correlation between predicted and observed frequencies of 0.68 in *S. cerevisiae* and 0.41 in *E. coli*, but only 0.16 in *M. tuberculosis*. While all three of these correlations are statistically significant (Table 1), it is clear that the predictive power of a model depending only on mutation rates and codon frequencies differs between these three species, an observation that we will return to shortly.

### Randomization tests support the relevance of empirical mutation spectra for adaptive evolution

The species-specific mutation spectra employed above reflect either (1) mutation-accumulation experiments under laboratory conditions in the absence of selection (*S. cerevisiae*, *E. coli*), or (2) the frequencies of putatively neutral single-nucleotide polymorphisms in natural populations (*M. tuberculosis*). We were struck by the observation that, using these spectra in a prediction model, the 95 % confidence interval on the mutation coefficient contained *β* = 1 for each of the three species. This observation not only suggests a strong influence of mutation bias on adaptation, but also that previously reported mutation spectra are relevant for adaptive evolution.

How well do these species-specific mutation spectra (reported in previously published studies) perform relative to randomly generated spectra, or to optimized spectra? To address this question, we repeated our analysis 10^6^ times, each time using a randomized mutation spectrum followed by the same negative binomial regression according to Eqn. 2. Each randomized spectrum was generated by drawing a random number between zero and one for each of the six possible mutation types, using a uniform distribution, and then normalizing the values by their sum to obtain a probability for each type. We then calculated the difference between the log-likelihood of the model fit with the randomized mutation spectrum and the log-likelihood of the model fit with the empirical mutation spectrum. When this difference is positive, the fit using the randomized mutation spectrum explains the spectrum of adaptive substitutions better than the fit using the empirical mutation spectrum, and when this difference is negative the empirical mutation spectrum provides the better explanation. Fig. 3a-c shows that the fit using the empirical mutation spectrum almost always explains the spectrum of adaptive substitutions better than fits using randomized mutation spectra, for all three species. Specifically, random mutation spectra outperformed empirical spectra with frequency 0.002 for *S. cerevisiae*, 0.037 for *E. coli*, and 0.035 for *M. tuberculosis*. This supports the hypothesis that the genetic changes favored by mutation are also those more likely to be used during adaptation, and highlights the relevance of empirically characterized mutation spectra for adaptive evolution in the laboratory (*S. cerevisiae*, *E. coli*) and in nature (*M. tuberculosis*).

**Fig. 3.**
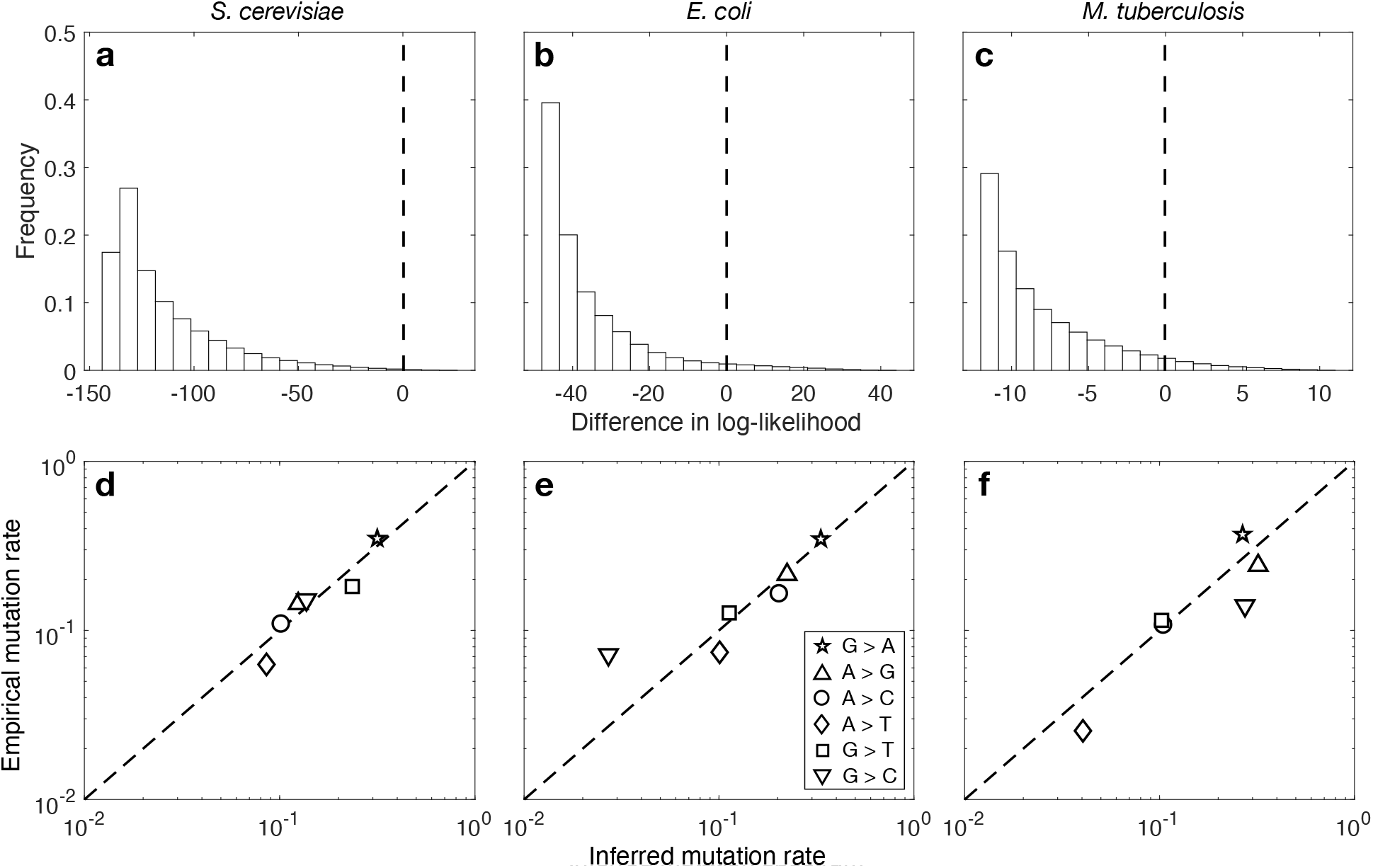
Empirical mutation rates explain the spectrum of adaptive substitutions better than randomized rates. In the upper panels, the white bars show the distribution of log-likelihood differences for randomized vs. empirical mutation rates for (a) *S. cerevisiae*, (b) *E. coli*, and (c) *M. tuberculosis*. A value of 0 (dashed vertical line) means that a simulated rate performs as well as the empirical mutation rate. The fraction of randomized rates providing a better model fit than the empirical rates (i.e., right of 0) is 0.2 %, 3.7 %, 3.5 % for panels a, b and c, respectively. Data based on 10^6^ randomized rates. Note that the three panels have different limits on their horizontal axes. In the lower panels, the empirical mutation rate is shown in relation to the inferred mutation rate on a double logarithmic scale for (d) *S. cerevisiae*, (e) *E. coli*, and (f) *M. tuberculosis*. Symbol types correspond to inset in (e). The dashed diagonal line indicates *y* = *x*.

While so far we have attempted to predict the spectrum of adaptive substitutions based on empirically observed mutation spectra, the strong relationship between the mutational and adaptive spectra in these three species suggests that it might also be possible to estimate the mutation spectrum from the spectrum of adaptive substitutions. To do this, we again fitted a negative binomial model but treated the rates of the six possible types of single nucleotide mutations as free parameters, which we estimated using maximum likelihood. We see that these inferred mutation spectra bear a strong resemblance to the experimentally characterized mutation spectra (Fig. 3d-f), with a Pearson correlation coefficient between the rates of 0.945 (*p* = 0.004) for *S. cerevisiae*, 0.960 (*p* = 0.002) for *E. coli*, and 0.827 (*p* = 0.042) for *M. tuberculosis*.

### What factors determine the predictive power of the model?

Although the analysis above reveals a statistically significant and approximately directly proportional contribution of mutational biases to the spectrum of adaptive substitutions for all three data sets, there is considerable variation in the strength of the correlation between the predicted and observed spectra, with this correlation being strongest and most significant for *S. cerevisiae*, and weakest and least significant for *M. tuberculosis* (Table 1).

One immediate hypothesis is that this variation in predictive power is driven by differences in the completeness of our estimates of the spectrum of adaptive substitutions. Even though our data sets include hundreds to thousands of adaptive events per species, a substantial fraction of the 354 possible types of codon-to-amino acid substitutions are missing from the spectrum for each species, a situation that likely arises both due to finite sample size effects and the limited diversity of distinct adaptive paths under a specific ecological circumstance (e.g., only a limited number of mutations confer resistance to any given antibiotic). Moreover, we note that at a qualitative level, the smaller the number of missing codon-to-amino acid paths, the stronger the correlation between predicted and observed spectra of adaptive substitutions (Table 1).

To better evaluate the influence of sparse sampling of codon-to-amino acid paths on the predictive power of our model, we simulated random data under our codon model with *β* = 1 (Eqn. 2), sampling adaptive events according to their expected frequencies, based on the empirical codon frequencies and mutation spectrum of each species, but restricting the sampled adaptive events to those corresponding to the non-zero elements of the observed spectra of adaptive substitutions. We then used negative binomial regression to fit this simulated spectrum of adaptive substitutions and measured the correlation between the randomized spectrum of adaptive substitutions and the spectrum of adaptive substitutions predicted by the fitted model. We repeated this process 10^3^ times for each species to obtain a distribution of correlations. Fig. S2 shows these distributions. On average, the correlations decreased from *S. cerevisiae* (0.76) to *E. coli* (0.75) to *M. tuberculosis* (0.61), suggesting that limitations in our data on the spectrum of adaptive substitutions are partly responsible for differences in model fits between the three species. However, Fig. S2 also shows that the correlations for these simulated data sets are considerably higher than those obtained with models fit to the observed spectra of adaptive substitutions (triangles in Fig. S2), suggesting the presence of other factors that modulate the predictive power of our modeling framework.

In order to address a combination of other potentially relevant factors, we turned to population-genetic simulations of evolution in a haploid genome, with variable parameters for population size *N*, mutation rate *μ*, and fraction of beneficial mutational paths *B*. The model genome consists of 500 codons subject to missense mutations, where a fraction *B* of such mutational paths are beneficial with a positive selection coefficient drawn from an exponential distribution, and all other paths are deleterious with effects drawn from a reflected gamma distribution (Methods). These simulations were implemented in SLiM v3.4 [32]. For each run of the simulation, we recorded the identity of the first adaptive mutation to reach fixation, repeating this process 1000 times to produce a simulated data set of adaptive substitutions of a similar size to our empirical data sets. For each of various combinations of *N*, *μ* and *B*, we then constructed 50 such simulated data sets (Methods) and analyzed these data sets using our negative binomial model.

Previous theoretical results suggest that the mutational supply (given by the product *Nμ*) should affect the extent to which mutational biases influence the distribution of adaptive substitutions [33–36]. In particular, the simplest effect of increasing *Nμ* is that multiple beneficial mutations are typically simultaneously present in the population, so that the adaptive mutation that ultimately fixes in the population is determined more by selective differences between these segregating mutations than by which beneficial mutation becomes established in the population first. Fig. 4a confirms the presence of this effect in our simulations by showing the inferred mutation coefficient *β* in relation to mutation supply (*Nμ*) for different proportions of beneficial mutations *B*. At the lowest mutation supply, *β* is approximately one, reflecting the direct proportionality between mutation rates and evolutionary outcomes that is expected in this regime [33, 37]. As the mutation supply increases, the average value of *β* tends toward zero, reflecting a diminished influence of mutation bias on adaptation. At the same time, the distribution of estimates for *β* becomes more dispersed (Fig. 4a) and the individual estimates become both less significant and less certain, as indicated by increasing average *p*-values and increasingly large confidence intervals (Fig. S3). Similarly, the predictive power of our model decreases with increasing mutation supply, as measured by a decreasing average correlation between the predicted and observed spectra of adaptive events (Fig. 4b).

**Fig. 4.**
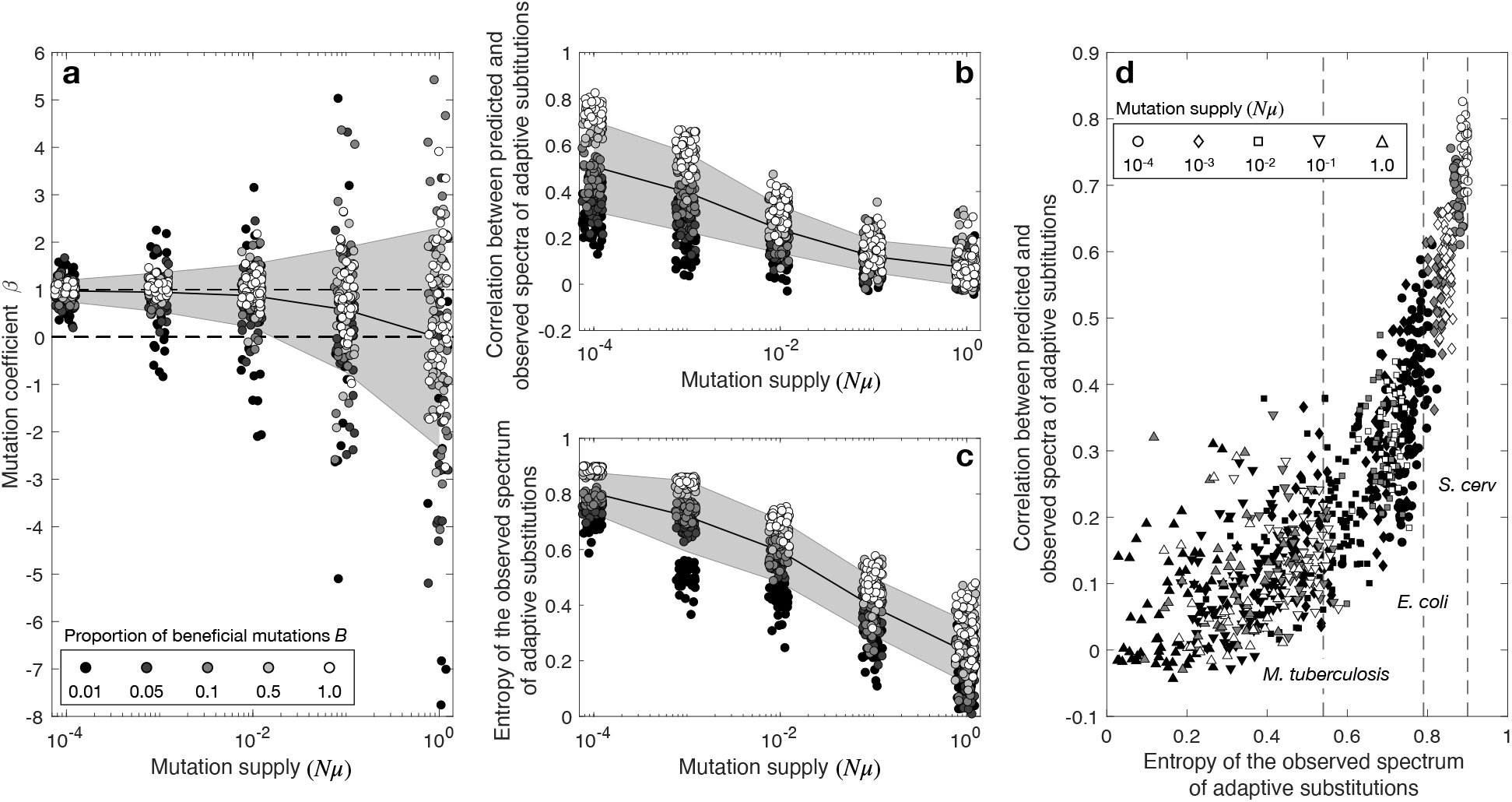
Evolutionary simulations show mutation supply and mutational target size jointly modulate the predictive power of our model. (a) The inferred mutation coefficient *β* as a function of *Nμ* for five different values of *B*, the fraction of beneficial mutations (the same color scheme for *B* is used in all panels). Dashed horizontal lines are drawn at *β* = 0 and *β* = 1 to indicate no influence and proportional influence of the mutation spectrum on the spectrum of adaptive substitutions, respectively. (b) Pearson’s correlation coefficient between predicted and simulated spectra of adaptive substitutions as a function of *Nμ* for five different values of *B*, and (c) entropy of simulated spectra of adaptive substitutions as a function of *Nμ* for five different values of *B*. In (a-c), the black lines show the mean and the gray areas show the standard deviation. (d) The Pearson’s correlation coefficient between predicted and simulated spectra of adaptive substitutions is shown in relation to the entropy of the simulated spectra of adaptive substitutions for different levels of mutation supply. The dashed vertical lines show the entropy of the spectrum of adaptive substitutions for each of our three study species.

The size of the mutational target also influences the predictive power of the fitted models, but in a somewhat more surprising manner. Intuitively, one might think that increasing the proportion of beneficial mutations would decrease the predictive power since this effectively increases the (beneficial) mutational supply, allowing increased competition between simultaneously segregating beneficial mutations. However, Fig. 4a and b show the opposite pattern, with low values of *B* showing the highest variability in estimated *β* values (Fig. 4a) and the lowest predictive power (Fig. 4b). We reason this occurs because larger mutational targets are more likely to contain a range of mutationally favored and disfavored paths in comparison to smaller mutational targets – thus allowing a correlation to emerge.

So far, we have shown that sparse sampling of codon-to-amino acid paths, increasing mutational supply, and a low proportion of beneficial mutations all tend to decrease the predictive power of our model. One unifying explanation for these observations rests on the fact that mutational biases have relatively broad effects on the spectrum of adaptive substitutions, in the sense that increasing a specific single-nucleotide mutation rate will cause a concomitant change in the relative frequencies of ∼60 distinct codon-to-amino acid paths. Thus, context-independent mutational biases result in the enrichment of broad classes of codon-to-amino acid substitutions and will therefore tend to perform poorly in predicting distributions of adaptive events that are highly concentrated on a small set of paths, whether this is because of relatively few available adaptive paths in a given selective environment (small *B*), limited sample size (zeros in observed spectrum), or a distribution of adaptive substitutions concentrated on the few fittest variants (large *Nμ*)

To quantify both the breadth of the adaptive spectrum and its effects on the predictive power of our model, we calculated the entropy of the spectrum of adaptive substitutions. We normalized the entropy so that it takes on its minimum value of 0 when all adaptive events correspond to a single codon-to-amino acid change and its maximum value of 1 when the adaptive events are uniformly distributed across all possible codon-to-amino acid changes (Methods). Thus, the entropy quantifies how evenly distributed the adaptive events are among the 354 possible codon-to-amino acid changes.

Fig. 4c shows that the entropy of the spectrum of adaptive substitutions indeed decreases as mutation supply increases, and that for any level of mutation supply, a lower proportion of beneficial mutations likewise decreases the entropy. To determine whether these patterns of decreasing entropy are sufficient to explain differences in the predictive power of our model across the range of model parameters, we plotted the correlation between predicted and observed events against the entropy of the spectrum of adaptive substitutions (Fig. 4d). We see that increasing entropy, either via a decreased mutation supply or an increased proportion of beneficial mutations, increases the correlation between simulated and predicted spectra of adaptive substitutions. These observations from the evolutionary simulations are qualitatively similar to our empirical observation that as the entropy of the spectrum of adaptive substitutions increases from *M. tuberculosis* to *E. coli* to *S. cerevisiae*, there is a corresponding increase in the correlation between predicted and observed spectra of adaptive substitutions (Table 1). Indeed the correlations for our three empirical data sets are well within the range of what we would expect from our simulations given their respective entropies (Fig. 4d). We thus conclude that many different factors could potentially influence the predictive power of our model via effects on the entropy of the spectrum of adaptive substitutions, and that these likely include both population genetic parameters such as mutation supply, as well as the genetic architecture of the trait being selected, and the number and diversity of adaptive challenges used to construct the spectrum of adaptive substitutions.

### Assessing possible effects of contamination

A key assumption of the analysis above is that the observations used to construct the spectrum of adaptive codon-to-amino acid changes are indeed adaptive. While this is likely the case for the *M. tuberculosis* data set, we now consider the possibility that some fraction of observations in the *S. cerevisiae* and *E. coli* data sets represent contamination such as hitchhikers. If contaminants reflect the mutation spectrum more than genuine adaptive changes, this will exaggerate the correspondence with mutational predictions.

Following [8], we use the observed dN/dS among all substitutions in the adapted lines to estimate the fraction of events in our data sets that are non-adaptive hitchhikers rather than adaptive drivers (Methods). We find such proportions to be ∼24% and ∼13% for *S. cerevisiae* and *E. coli*, respectively. We then assess the influence of contamination by randomly removing a fraction *q* of observations, sampled according to the empirical mutation spectrum: this procedure simulates the removal of a hypothetical contaminant fraction of size *q* under the worst-case scenario that the nucleotide changes in the contaminant fraction mirror the mutation spectrum. As shown in Fig. S4, even under the assumption that 40% of the mutations are contaminants, we observe a strong and statistically significant influence of mutation bias on adaptive evolution. In fact, we estimate that ∼65% and ∼44% of contamination—for *S. cerevisiae* and *E. coli*, respectively—would be required to increase the *p*-value of *β* to the point where the influence of mutation bias would no longer be detectable.

We only carried out this procedure for the *S. cerevisiae* and *E. coli* data sets, because they include all missense changes in the genomes of adapted strains, rather than only driver mutations that are verified experimentally, and are therefore likely to include a minority of hitchhikers [28, 38]. By contrast, the *M. tuberculosis* data set only includes mutations that have been shown experimentally to confer antibiotic resistance [5]. This kind of data set represents the ideal that, perhaps, can be expected to predominate in the future, as it becomes easier to carry out genome editing and functional assays in a high-throughput manner.

## DISCUSSION

A growing body of evidence suggests that specific mutation biases influence the types of genetic changes that cause adaptation [5, 19–26], consistent with a small body of theoretical work on how biases in the introduction of variation— both low-level mutational biases and higher-level systemic biases—are expected to influence evolution [33–36]. Here, we have developed and applied a general approach to assess how the mutation spectrum shapes the spectrum of adaptive substitutions. Our approach uses negative binomial regression to model the spectrum of adaptive substitutions as a function of codon frequencies and the mutation spectrum, measuring the influence of mutation in terms of the regression coefficient *β*. Such an approach can be applied to any sufficiently large data set of substitutions associated with adaptation, given codon frequencies and an estimate of the mutation spectrum. Applying our model to three such species (*Saccharomyces cerevisiae*, *Escherichia coli*, and *Mycobacterium tuberculosis*), we uncovered a clear signal that the mutation spectrum shaped the spectrum of adaptive substitutions. The influence of mutation bias on the spectrum of adaptive substitutions is proportional in the sense that the inferred value of *β* is not significantly different from 1 in any species. This result holds even when we account for contamination by hitchikers in the data sets for *S. cerevisiae* and *E. coli*.

Our approach also illustrates how the spectrum of adaptive substitutions may be interrogated to reveal clues about the genetic basis of adaptation. We used our fitted models to predict the spectrum of adaptive substitutions in each species, and uncovered variation in their predictive capacity, decreasing from *S. cerevisiae* to *E. coli* to *M. tuberculosis*. Using evolutionary simulations, we uncovered multiple potential sources of this variation. Specifically, we found that the degree to which the mutation spectrum is a good predictor of the spectrum of adaptive substitutions depends on how the adaptive events are distributed amongst all possible codon-to-amino acid changes, with distributions concentrated on a small number of codon-to-amino acid changes associated with reduced predictive capacity. Factors that affect this distribution include data set size, population genetic conditions, diversity of selective environments, and the genetic architecture of adaptive traits. Importantly, population genetic conditions that modulate the influence of mutation bias on adaptation, such as mutation supply, and non-population genetic conditions, such as the diversity of environmental conditions included in the data set, can affect the predictive capacity of our model in similar ways. Additional work is needed to disambiguate these various causes of differing model fits between species.

For example, the three species studied here vary in their population genetic and environmental conditions, as well as their mutational target sizes. *M. tuberculosis* has one of the lowest mutation supplies of all bacteria [39], a small population size upon infection [40], and the 11 antibiotics considered here target specific gene products [5]. For example, Rifampicin targets the beta subunit of bacterial RNA polymerase, and only a small handful of mutations to the *rpoB* gene that encodes this subunit cause resistance [41]. Thus, while the population genetic conditions of *M. tuberculosis* are more likely similar to origin-fixation dynamics than clonal interference dynamics, and the set of observations is large, the mutational target size for antibiotic resistance is small. In contrast, *E. coli* experiences clonal interference due to a relatively higher mutation supply [38], but adaptation to temperature stress involves a larger mutational target [8, 42]. Similarly, *S. cerevisiae* experiences clonal interference due to a high mutation supply [28], but because the data we study include adaptation to several environmental conditions, the mutational target size is large. Thus, the inferred influence of mutation bias on adaptation in these three species, increasing from *M. tuberculosis* to *E. coli* to *S. cerevisiae*, is consistent with our findings from evolutionary simulations that mutation supply and mutational target size modulate the influence of mutation bias on adaptation.

Though this simple model has proven useful, further work may benefit from a broader consideration of sources of heterogeneity. For instance, a more sophisticated treatment of the mutation spectrum would include effects of local sequence context [43, 44]. Likewise, the influence of the genetic code could be parameterized separately, as a step toward understanding the broader evolutionary issue of how genotype-phenotype maps shape the course of evolution.

Our analysis of mutational effects includes heterogeneity in fitness effects among beneficial paths (captured implicitly via the dispersion parameter of the negative binomial model), but does not suppose any systematic relationship between fitness and codon-to-amino-acid paths. If some beneficial codon-to-amino acid changes have systematically higher selection coefficients than others, which one might expect from generic differences in amino acid exchangeability [45], this may influence how strongly the mutation spectrum shapes the spectrum of adaptive substitutions. If the nucleotide changes favored by mutation are not the same as those favored by selection, this could diminish the influence of mutation bias on adaptation [35]. This kind of effect might be particularly strong due to the dominance of a small number of idiosyncratic paths. That is, if fitness effects are highly heterogeneous, such that a small number of mutations have exceedingly high selection coefficients, and these nucleotide changes are not those favored by mutation, this could diminish the predictive capacity of our model. The data set for *M. tuberculosis* contains such “jackpot” mutations [5], e.g., the G→C transversion that causes the S315T substitution in KatG and confers resistance to isoniazid [30]. Because the mutation spectrum of *M. tuberculosis* is biased toward transitions [12], this jackpot mutation likely reduces the predictive capacity of our fitted model.

The discovery that the mutation spectrum strongly shapes the spectrum of adaptive substitutions has several implications. First, this finding has implications for the predictability of evolution [46–48], because it shows that the nucleotide changes that are more likely to arise via mutation are also those more likely to contribute to evolutionary adaptation, an effect that is both large and readily predictable from data on the mutation spectrum. Long-term laboratory evolution experiments often uncover molecular diversity in adaptive convergence, meaning that in replicate populations, distinct sets of mutations cause adaptation to identical environments [8]. We uncover an additional layer of convergence: though distinct sets of mutations cause adaptation in different replicate populations, the influence of mutation bias causes these sets to converge on similar patterns of nucleotide changes and codon-to-amino-acid changes.

Secondly, the discovery of a direct influence of mutation bias on evolutionary adaptation parallels recent reports that driver mutations in cancer reflect the underlying biases of cancer-associated mutational processes, including exogenous effects of UV light and tobacco exposure, and endogenous effects of DNA mismatch repair and APOBEC activity [49–51]. The increased predictability of such changes, due to mutational effects, can inform rational drug design, as has been suggested for drugs for leukemia, prostate cancer, breast cancer, and gastrointestinal stromal tumors [26]. The same may be true for designing antibiotic treatments for mycobacteria, which evolve multi-drug resistance via a sequence of mutations, several of which interact epistatically, such that only a subset of possible mutational trajectories to multi-drug resistance are possible [52]. If some of these paths comprise nucleotide changes that are less likely to arise via mutation, then this could inform treatment regimens.

Finally, the broadest context for the present work is a debate about the relative roles of mutation and selection in shaping the course of evolution. Arguments dating back to the Modern Synthesis emphasize selection as the sole directional force, with mutation treated as a weak and ineffectual pressure due to the smallness of mutation rates [53–55], e.g., Haldane concluded that mutation can influence the course of evolution only under neutral evolution, or when mutation rates are unusually high [53]. More recent theory shows how such conclusions depend on assuming that evolution begins with abundant standing genetic variation, so that mutation acts only as a frequency-shifting force and not as the source of genetic novelty [33]. When evolution depends on mutation as a source of novelty, biases in the introduction of variants, such as toward particular nucleotide changes, systematically influence which genetic changes are involved in adaptation [34, 56].

Some authors have responded to the theory of mutation-biased adaptation by arguing that such an influence is unlikely, on the grounds of requiring sign epistasis or unusually small population sizes [57]. However, modeling here and in other work [35, 36] shows that mutation bias can influence adaptation across a range of conditions, including conditions that induce clonal interference among concurrent mutations. More broadly, while theoretical arguments are surely helpful for sharpening our understanding, ultimately the prevalence and magnitude of the mutational influence on adaptation is an empirical question, and the impact of mutational biases has now been shown for several different types of mutations, in a range of systems from bacteriophage to birds to somatic evolution in human cancers [5, 19–26].

This growing body of work, in turn, provides a population-genetic mechanism for previously proposed theories concerning how variational properties influence the evolutionary process. For instance, evo-devo arguments about bias or constraint relate evolutionary patterns to tendencies of developmental variation, but the causal nature of this link, in terms of population-genetic principles, is typically unspecified (e.g., [58,59]). Though some sources invoke constraints in the context of quantitative genetics [60], the latter framework only applies to dimensional biases in quantitative traits, whereas the theory of biases in the introduction process is suitable for molecular and other discrete traits, e.g., this theory plausibly applies to a small body of work on the tendency of evolution to prefer more findable structures in cases such as RNA folds [61] or regulatory circuits [62]. Our results improve the population-genetic underpinnings of these theories by showing that mutational biases, which are a similar but even simpler set of biases, have a clear and measurable impact on the distribution of variants fixed during adaptive evolution.

## METHODS

### Data

Our modeling framework is built around three key quantities, which are specific to each species: A spectrum of adaptive substitutions **n**, a table of codon frequencies *f*, and a mutation spectrum *μ*. These are all constructed using empirical data, as described below.

#### Spectrum of adaptive substitutions

We curated a list of missense mutations associated with adaptation from the published literature for each of three species: *S. cerevisiae*, *E. coli*, and *M. tuberculosis*. For each mutation, these lists specify a genomic coordinate, nucleotide change, amino acid substitution, and literature reference (Tables S2-S4). We refer to each unique combination of genomic coordinate and nucleotide change as a mutational path and each instance of adaptive change along a mutational path as an adaptive event. The number of adaptive events per mutational path are also reported in Tables S2-S4.

For *S. cerevisiae*, the adaptive events were reported in four studies, each of which considered one or more environmental or genetic challenges, including high salinity [27], low glucose [27], rich media [28], and gene knockout [29]. The list contains 721 adaptive events across 534 mutational paths (Table S2).

For *E. coli*, the adaptive events were reported in a single study of 115 replicate populations adapting to temperature stress [8]. The list contains 602 adaptive events across 492 mutational paths (Table S3).

For *M. tuberculosis*, the adaptive events were reported in a single study of the influence of mutation bias on adaptation to antibiotic stress [5]. The underlying mutational paths were derived from two separate meta-analysis of the literature on antibiotic resistance (one performed for the study and another previously published [4]), with each mutational path required to pass stringent tests for conferring antibiotic resistance. A total of 11 antibiotics or antibiotic classes were considered: Rifampicin, ethambutol, isoniazid, ethionamide, ofloxacin, pyrazinamide, streptomycin, kanamycin, pyrazinamide, fluoroquinolones, and aminoglycosides. The adaptive events were inferred from a phylogenetic reconstruction of public *M. tuberculosis* genomes. We merged the adaptive events from the two meta-analyses. The resulting list contains 4413 adaptive events across 283 mutational paths (Table S4). Analyzing the adaptive events from the two meta-analyses separately (Table S1) produced qualitatively similar results to those reported in Table 1.

For each species, we constructed the spectrum of adaptive substitutions **n** from the list of adaptive events described above, assigning each adaptive event to its respective codon-to-amino-acid change. Each element **n**(*c, a*) of the spectrum of adaptive substitutions therefore tallies the number of adaptive events that changed codon *c* to amino acid *a*. Note the adaptive events tallied for any codon-to-amino-acid change often reflect more than one genomic coordinate and/or nucleotide change (i.e., different mutation paths). These spectra are reported in Table S5.

#### Codon frequencies

We used the tables of codon frequencies reported in the Codon Usage Database [63], found via query to an exact match to *Saccharomyces cerevisiae*, *Escherichia coli*, and *Mycobacterium tuberculosis*. These frequencies are reported in Table S6 and shown in Fig. S1e-g.

#### Empirical mutation spectra

For *S. cerevisiae* and *E. coli*, we used mutation rates derived from mutation accumulation experiments, as reported in Figure 3 of reference [15] and Table 3 of reference [14], respectively. For *M. tuberculosis*, we used mutation rates derived from single-nucleotide polymorphism data, extracted from Figure 2A in reference [12] using a web-based image analysis tool [64]. For *E. coli*, we corrected the mutation rates for GC content, following [12]. For *S. cerevisiae* and *M. tuberculosis*, the rates were already corrected [12, 15].

These spectra are reported in Table S7 and shown in Fig. S1a. We used these estimated mutation rates to define a total codon-to-amino acid mutation rate *μ*(*c, a*) for each of the 354 codon-to-amino acid changes allowed by the standard genetic code, summing the rates of all point mutations in codon *c* that lead to amino acid *a*. For example, the probability of the substitution from codon CAC to Glutamine (Q) is the sum of the probabilities of point mutations C→A and C→G, since both mutations in the third position of CAC lead to codons for Glutamine (Q).

### Entropy of the spectrum of adaptive substitutions

The spectrum of adaptive substitutions **n** describes the number of adaptive events per codon-to-amino acid change. We calculate the entropy *H* of this spectrum as

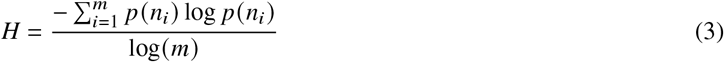

where *p* (*n_i_*) is the proportion of adaptive events that correspond to the *i*th codon-amino acid change, and *m* = 354 is the number of codon-to-amino acid changes allowed by the standard genetic code.

### Evolutionary simulations

We used SLiM v3.4 for the evolutionary simulations [32]. We ran each simulation until a single mutation went to fixation, which we recorded as an adaptive event. We recorded 1000 such events per replicate by running 1000 independent simulations. We performed 50 replicates per combination of the parameters *N*, *μ*, and *B*.

Each of the 1000 simulations per replicate used the same initial population, which comprised *N* copies of a nucleotide sequence of length *L* = 1500 (i.e., 500 codons), randomly generated using the codon frequencies for *S. cerevisiae*. All sequences in the initial population were assigned a fitness of one. The fitness effects assigned to each of the possible codon-to-amino acid changes from each of the 500 codons were drawn at random from a distribution of fitness effects, and were held constant across the 1000 simulations per replicate.

A unique distribution of fitness effects was constructed for each replicate, such that synonymous mutations were neutral, a fraction *B* of non-synonymous codon-to-amino acid changes were beneficial, and a fraction 1 − *B* of non-synonymous codon-to-amino acid changes were deleterious. The fitness effects of beneficial codon-to-amino acid changes were drawn from an exponential distribution with density

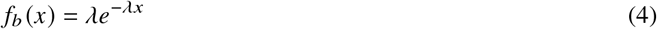

where *λ* = 33.33, so that the expected advantageous selection coefficient was 0.03. The fitness effects of deleterious codon-to-amino acid changes were drawn from a gamma distribution with density

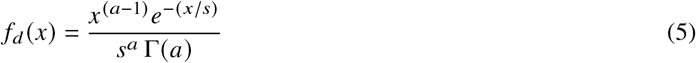

where *a* = 0.2 and *s* = 6.6. Fig. S5 shows representative distributions of fitness effects for different proportions of beneficial mutations *B*.

Each simulation proceeded until a single mutation went to fixation. In each generation *t*, *N* sequences were chosen from the population at generation *t* − 1 with replacement and with a probability proportional to their fitness. Mutations were introduced according to the product of the genome-wide mutation rate *μ* and the per-nucleotide mutation rate defined by the mutation spectrum for *S. cerevisiae*, with each mutation affecting fitness as defined at the onset of the simulation.

### Contamination estimates

For each type of mutation, we calculated the number of synonymous and non-synonymous sites for each possible codon, and we estimated the total number of synonymous and non-synonymous sites in the genome by taking into account the codon usage patterns of *S. cerevisiae* and *E. coli* (Fig. S1e-f). We then calculated dN/dS ratios among all substitutions in the adapted lines correcting for the mutation rates of each type of mutation (Fig. S1a). Following [8], we estimated the proportion of adaptive non-synonymous mutations from such ratios as *y* = (*x* − 1.0)/*x*, where *x* is the estimated dN/dS ratio (4.24 and 7.76 for *S. cerevisiae* and *E. coli*, respectively). Finally, we estimated the fraction of hitch-hikers in our data sets as 1 − *y*.

## ACKNOWLEDGMENTS

The identification of any specific commercial products is for the purpose of specifying a protocol, and does not imply a recommendation or endorsement by the National Institute of Standards and Technology. This project / publication was made possible through the support of a grant from the John Templeton Foundation (grant #61782, D.M.M.) and from the Swiss National Science Foundation (grant #PP00P3_170604, J.L.P.). The opinions expressed in this publication are those of the authors and do not necessarily reflect the views of the John Templeton Foundation. D.M.M. also acknowledges additional support from an an Alfred P. Sloan Research Fellowship and from the Simons Center for Quantitative Biology.

---

**TABLE S1.** Separately analyzing the adaptive events from the two meta-analyses of antibiotic resistance mutations in *M. tuberculosis* yields qualitatively similar results to analyzing them together. Shown are the observed numbers of paths and events, the mutation coefficient *β* (with standard error) and its *p*-value, the Pearson’s correlation between observed and predicted spectra of adaptive substitutions and its *p*-value, as well as the number of non-zero elements of the spectrum of adaptive substitutions and the entropy of the spectrum of adaptive substitutions.

|  | Data |  | Neg. binomial regression |  | Prediction model |  | Spectrum elements |  |
| --- | --- | --- | --- | --- | --- | --- | --- | --- |
| Study | Paths | Events | $\beta$ | $p_\beta$ | Correlation | $p_{\text{corr}}$ | Non-zero elements | Entropy |
| Basel [5] | 126 | 2319 | $0.86 \pm 0.27$ | 0.001 | 0.15 | 0.005 | 78 | 0.53 |
| Manson [4] | 168 | 2094 | $0.86 \pm 0.27$ | 0.002 | 0.17 | 0.002 | 80 | 0.52 |

**Fig. S1.**
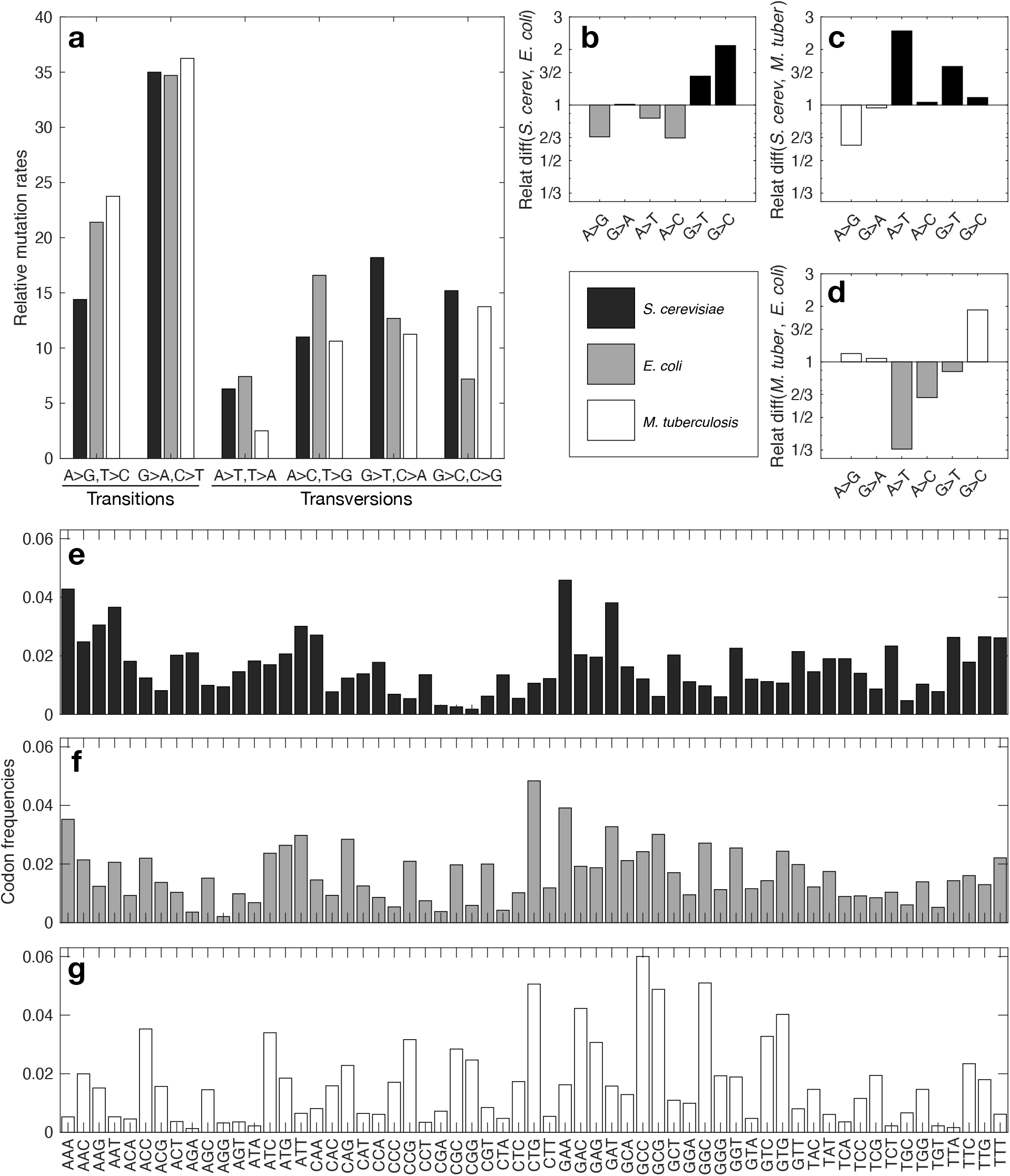
Empirical mutation spectra and codon frequencies. (a) Bar plots of the empirical mutation spectra for *S. cerevisiae*, *E. coli*, and *M. tuberculosis*. Bar color indicates the species; see legend. (b-d) Relative difference in mutation rates per mutation type, Relat diff (*b, a*) = *b/a*. Bar color indicates the species with the highest mutation rate for each mutation type. The vertical axis is logarithmically scaled for visual clarity. (e-g) Bar plots of the empirical codon frequencies for (e) *S. cerevisiae*, (f) *E. coli*, and (g) *M. tuberculosis*.

**Fig. S2.**
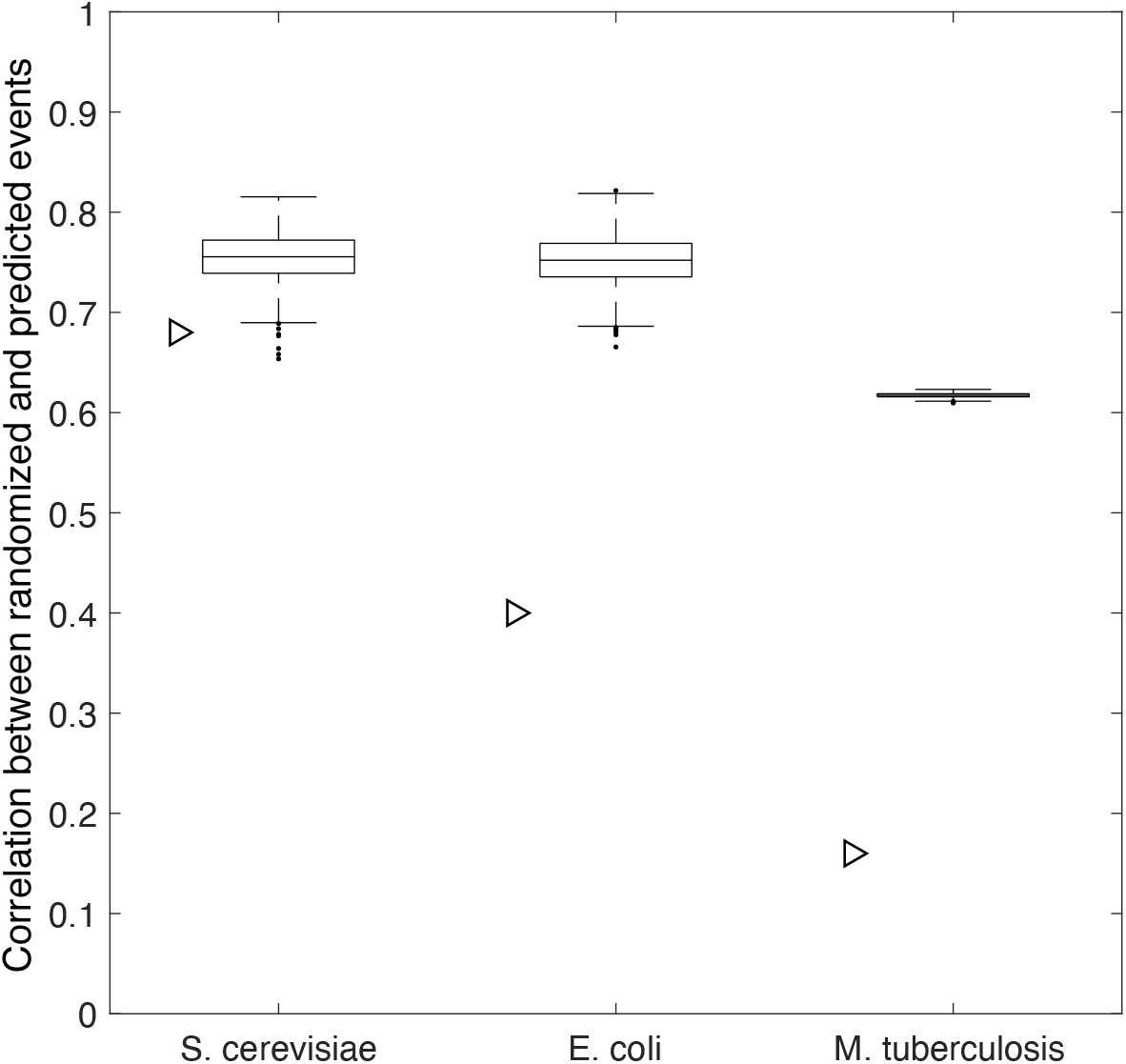
The correlation between predicted and randomized spectra of adaptive substitutions depends on mutational target size, even under origin-fixation dynamics. The distribution of correlations between predicted and randomized spectra of adaptive substitutions using the codon frequencies, mutation spectra, and number of non-zero elements in the spectrum of adaptive substitutions are shown for *S. cerevisiae*, *E. coli*, and *M. tuberculosis*. Data pertain to 10^3^ simulations. Triangles show the correlations reported in Table 1, for reference.

**Fig. S3.**
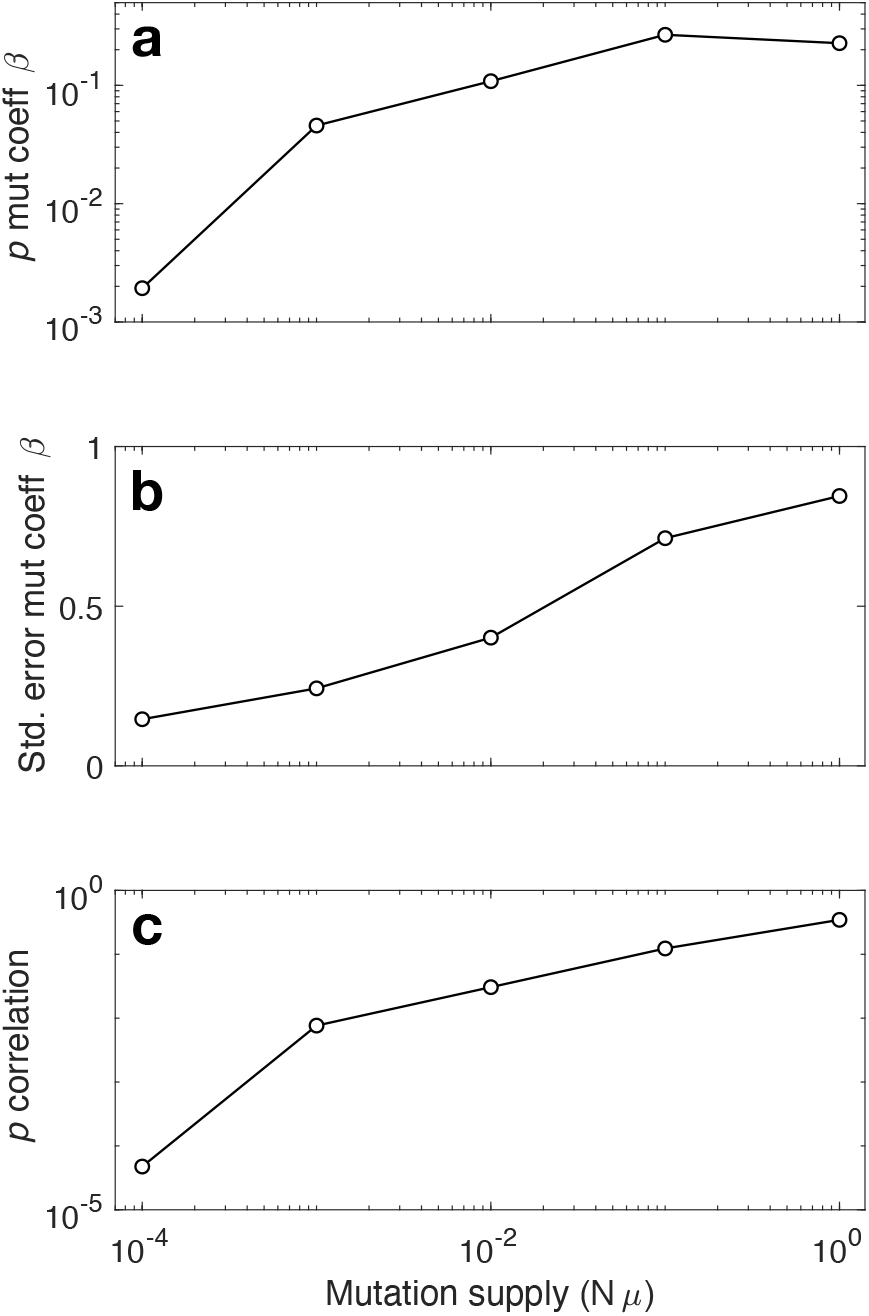
High mutation supply diminishes the influence of mutation bias on adaptive evolution. The a) average *p*-value and b) standard error of the mutation coefficient *β*, and c) the average *p*-value of the correlation between predicted and simulated spectra of adaptive substitutions are shown in relation to mutation supply *Nμ*. Data pertain to those shown in Figs. 4a-c.

**Fig. S4.**
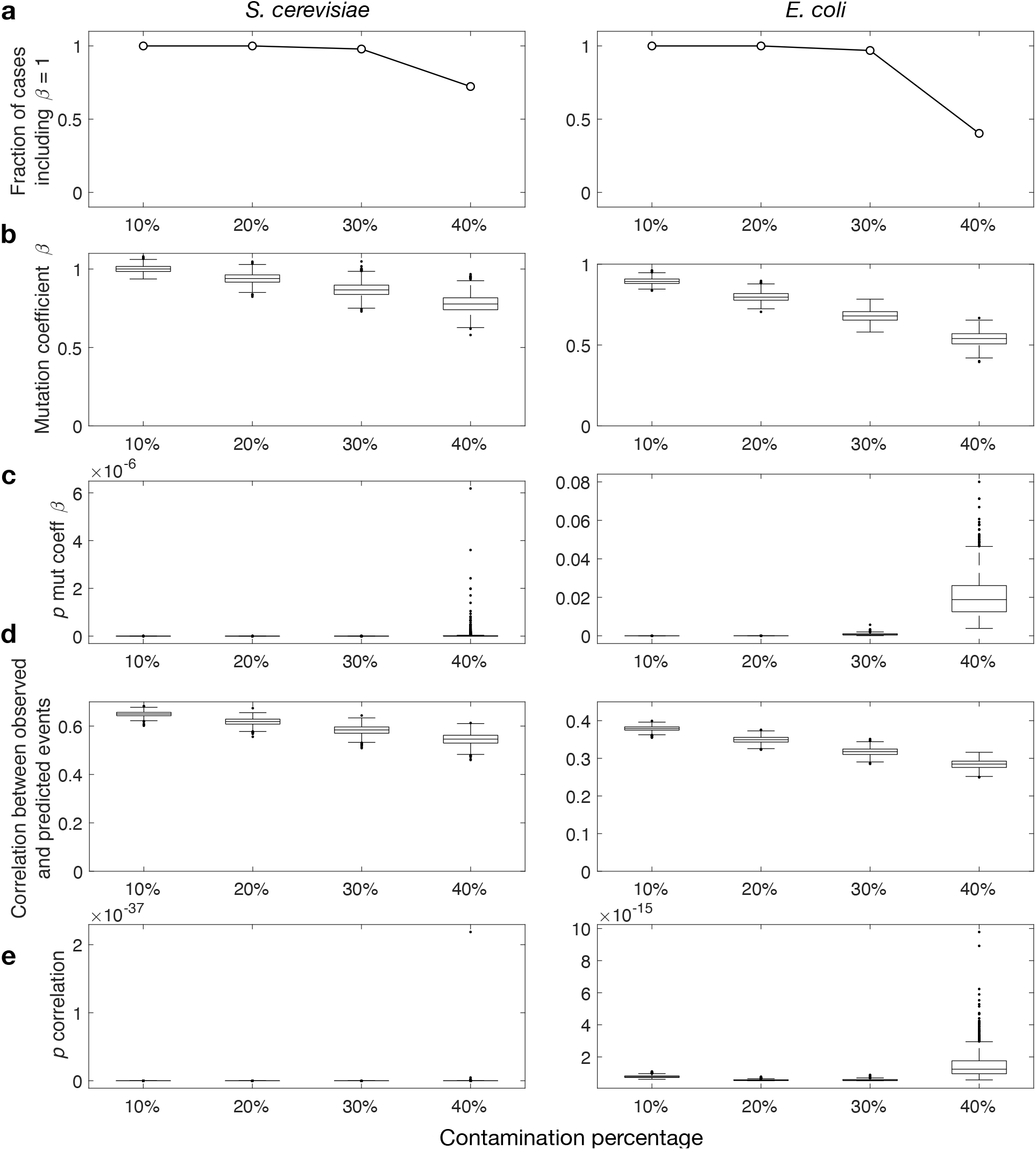
Contamination analysis supports the influence of mutation bias on adaptation. (a) Fraction of simulated data sets in which the confidence interval includes *β* = 1. (b) Inferred mutation coefficients*β*, (c) *p*-values of the regression coefficients *β*, (d) Pearson’s correlation coefficients between observed and predicted spectra of adaptive substitutions, and (e) the *p*-values of the correlation coefficients, are all shown in relation to the percentage of mutations randomly removed from the data sets of adaptive mutations.

**Fig. S5.**
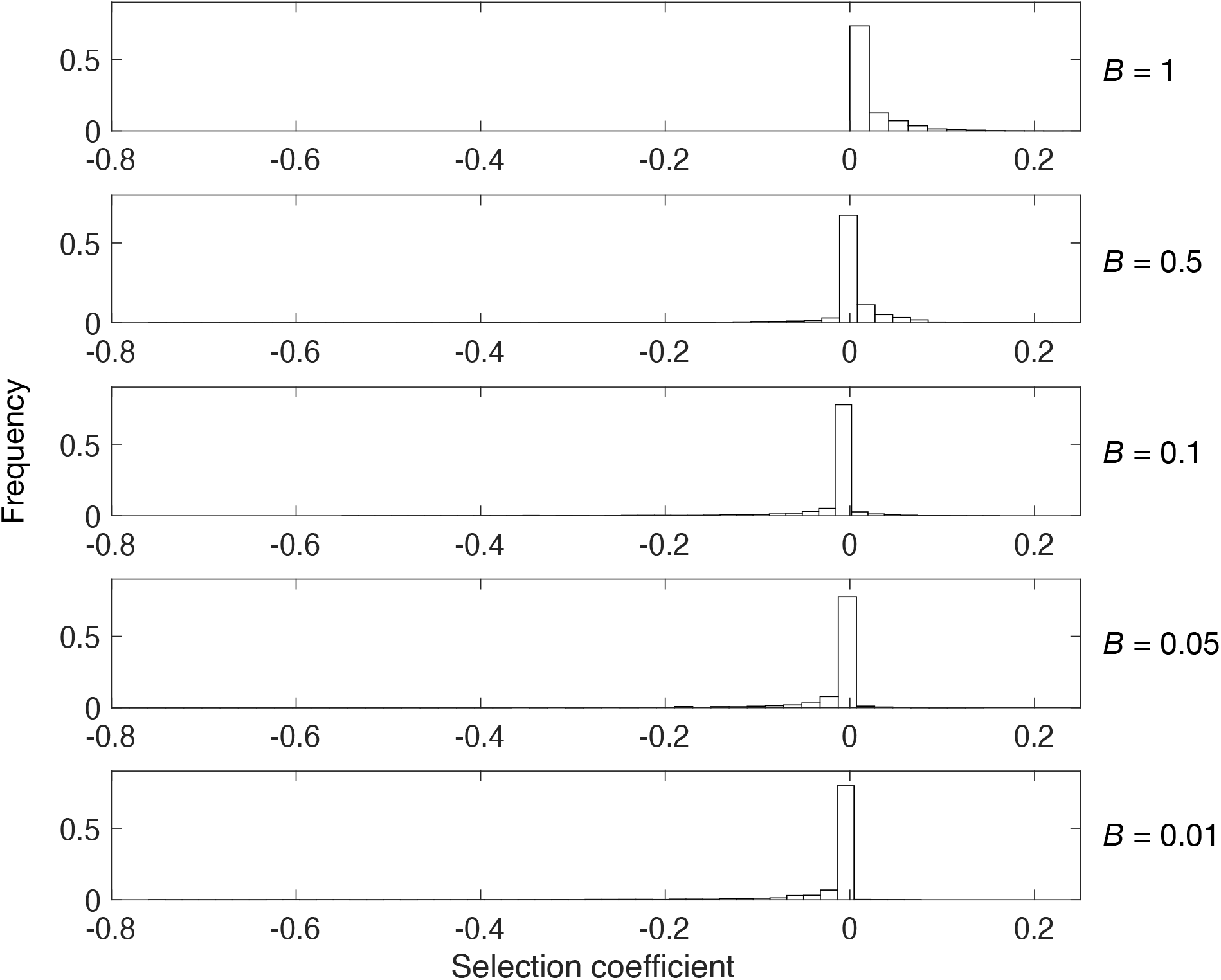
Distributions of fitness effects. Representative distributions of fitness effects used in the evolutionary simulations for five different proportions of beneficial mutations *B*.

